# Using molecular ecological network analysis to explore the effects of chemotherapy on intestinal microbial communities of colorectal cancer patients

**DOI:** 10.1101/331876

**Authors:** Jing Cong, Jingjuan Zhu, Chuantao Zhang, Tianjun Li, Kewei Liu, Dong Liu, Na Zhou, Man Jiang, Helei Hou, Xiaochun Zhang

## Abstract

Intestinal microbiota is now widely known to be key roles in the nutrition uptake, metabolism, and the regulation of human immune responses. However, we do not know how changes the intestinal microbiota in response to the chemotherapy. In this study, we used network-based analytical approaches to explore the effects of five stages of chemotherapy on the intestinal microbiota of colorectal cancer patients. The results showed that chemotherapy greatly reduced the alpha diversity and changed the specie-specie interaction networks of intestinal microbiota, proved by the network size, network connectivity and modularity. The OTU167 and OTU8 from the genus *Fusobacterium* and *Bacteroides* were identified as keystone taxa by molecular ecological networks in the first two stages of chemotherapy, and were significantly correlated with tumor makers (*P* < 0.05). Five stages of chemotherapy did not make the intestinal micro-ecosystem regain a steady state, because of the lower alpha diversity and more complicated ecological networks compared to the healthy individuals. Furthermore, combing the changes of ecological networks with the tumor markers, the intestinal microbiota was closely linked with clinical chemotherapeutic effects.

**Importance:** A deeply understanding of the role of intestinal microbiota contributes to help us find path forward for improving the prognosis of colorectal cancer patients. In addition, diet or probiotics interventions will be a possible attempt to improve the clinical chemotherapeutic effects for colorectal cancer patients.

## Introduction

Colorectal cancer (CRC) is a major killer of people in the world all the time, though continuing medical advances on several fronts. The CRC etiology is very complicated and still not fully understood, but CRC carcinogenesis is a heterogeneous progression with a train of genetic and epigenetic variations, and is profoundly influenced by dietary, environment, host and intestinal microbiota (1). Although genetic predisposition is closely associated with certain types of CRC, intestinal microbiota increasingly plays an indispensable role in human intestinal health. It has been proved that intestinal microbiota is concerned with the initiation and progression of colorectal cancer (2, 3). Heterogeneity of CRC-associated microbiota might reflect discrepancies in the host immunoinflammatory (4). Despite such heterogeneity, considering a combination of several bacteria in the faecal microbiota of CRC individuals as a marker for detecting the disease is feasible (5, 6). As a consequence, the plasticity of intestinal microbiota can be leveraged for therapeutic interventions (7) and to improve therapeutic effect (8).

Cytotoxic drugs continue to be the mainstay of therapy for most CRC patients, whereas the related treatment response is unpredictable. The personalized cancer therapies are now emerging, and targeted therapies have made revolutionized outcomes in CRC (9). However, it still appears novel problems such as idiosyncratic adverse effects, acquired resistance and high costs (10, 11). Recent studies have implicated intestinal microbiota at the species level in influencing the drug response and toxicity of CRC patients (12). Drug metabolism by intestinal microbiota was well recognized since the 1960s (13). Intestinal microbiota played a key role in confirming the efficacy and toxicity to a broad range of drugs (14). With the development of high-throughput sequencing, the importance of intestinal microbiota for drug modulation and discovery is increasingly recognized (15). Timothy A. Scott et al. reported that intestinal microbiota had an influence on fluoropyrimidines, which are the first-line treatment for CRC, through drug interconversion involving bacterial vitamin B6, B9, and ribonucleotide metabolism (16). Leah Guthrie et al. suggested that metagenomic mining of the microbiome, associated with metabolomics, was considered as a non-invasive approach to develop biomarkers for CRC treatment outcomes (2). These findings highlight intestinal microbiota as the potential therapeutic power to ensure host metabolic health and disease treatment.

Microbes coexist in complex environments in which interactions among individuals are indispensable for community assembly and ecosystem function (17, 18). However, few studies explore the interactions among intestinal microbiota, or determine which individuals share niches within the intestinal environment. Therefore, identifying and defining the interactions that occur among intestinal microbiota contributes to understanding microbial diversity and function to provide a possibility in better therapeutic interventions. Molecular ecological network analysis provides a promising future for exploring the dynamics of microbial interactions and niches (19).

Network analysis can identify putative keystone taxa which are important in maintaining community structure and function (20). In recent years, molecular ecological network analysis has been used as a tool to determine complex microbial assemblages in various environments such as human (21), groundwater (22) and soil (23).

To identify intestinal microbial assemblages that potentially interact or share niches within intestine, we used molecular ecological network analysis (24) to construct co-occurrence networks for healthy volunteers and CRC patients throughout the treatment. We conducted a study of intestinal microbiota using faecal samples from CRC patients and healthy volunteers. High-throughput sequencing of 16S rRNA gene amplicons was used to describe the intestinal bacterial assemblages. We mainly examined the changes of intestinal microbiota during the treatment based on the molecular ecological network. Molecular network analysis was used to address three questions: (1) if the microbial diversity and community structure were affected by chemotherapy treatment for CRC patients? (2) How does the bacterial interaction change during the treatment for CRC patients? (3) Are there taxa that play particularly important roles within CRC networks, suggesting that they may serve as keystone taxa in CRC communities? Our work identifies a previously undocumented dimension of the intestinal microbiota, and offers insight into the specie-specie interaction networks of intestinal microbiota associated with CRC patients in response to the chemotherapy.

## Results

### Variations of alpha diversity of intestinal microbiota from CRC patients in response to the chemotherapy

To explore the changes of intestinal microbiota for CRC patients in response to the chemotherapy, we examined the alpha diversity (Shannon diversity, Simpson diversity, phylogenetic diversity and richness) of intestinal microbiota (Table S1). The results showed that T1 had the lowest of Shannon diversity, phylogenetic diversity and richness, except for the Simpson diversity. The alpha diversity of intestinal microbiota in CRC patients was distinctly lower compared with the healthy individuals (Table S1). Based on the relative abundance, *Bacteroidaceae*, *Ruminococcaceae* and *Lachnospiraceae* were the most abundant family in healthy individuals, followed by *Prevotellaceae* and *Veillonellaceae* (Figure S1). In the CRC patients, except for *Bacteroidaceae*, *Ruminococcaceae* and *Lachnospiraceae* as main family, *Enterobacteriaceae* became more conspicuous in response to the chemotherapy. Some members of the *Enterobacteriaceae* produce endotoxins that cause a systemic inflammatory response when released into the bloodstream following cell lysis.

### Characteristics of constructed networks

To identify potential intestinal microbe-microbe interactions and niche-sharing in CRC patients’ intestine, we constructed microbial co-occurrence networks and further measured their topological parameters during abnormal microbial succession over treatment process (Figure 1 and Table 1). The identical threshold values imposed 0.66 in H and T samples. All the constructed networks showed scale-free characteristics, which means that only a few nodes in the network have a great many of connections, whereas most nodes have no or few connections (25), as proved by R^2^ of power law ranging from 0.72 to 0.87 (Table 1). Less modularity values (0.40-0.50) in T samples indicated that these networks could be impossibly isolated into multiple modules. The networks from the T0 to the T5 presented inconspicuous regularity, identified by the changes of the average connectivity (*avgK*), harmonic geodesic distance (*HD*) and modularity (Table 1). In addition, the constructed networks were significantly different from random networks obtained using identical numbers of nodes and links (*P* < 0.05, Tables 2). These metrics suggested that the network structures were non-random and unlikely due to chance.

**Table 1.** Topological properties of the ecological networks of intestinal microbiota from healthy individuals and colorectal cancer patients during the different treatment stages

|  | Sample | H | T0 | T1 | T2 | T3 | T4 | T5 |
| --- | --- | --- | --- | --- | --- | --- | --- | --- |
| Empirical<br>Network | Total nodes | 105 | 99 | 102 | 103 | 80 | 103 | 103 |
|  | Total links | 110 | 238 | 228 | 239 | 173 | 210 | 231 |
|  | Average degree<br>(avgK) | 2.10 | 4.81 | 4.47 | 4.64 | 4.33 | 4.08 | 4.49 |
|  | Average clustering<br>coefficient (avgCC) | 0.07 | 0.19 | 0.21 | 0.18 | 0.14 | 0.12 | 0.24 |
|  | Harmonic geodesic<br>distance (HD) | 4.17 | 2.49 | 2.54 | 2.76 | 2.72 | 3.02 | 2.67 |
|  | Identical threshold | 0.66 | 0.66 | 0.66 | 0.66 | 0.66 | 0.66 | 0.66 |
|  | R<sup>2</sup> of power-law | 0.79 | 0.79 | 0.72 | 0.82 | 0.81 | 0.86 | 0.87 |
|  | Modularity | 0.77 | 0.43 | 0.45 | 0.48 | 0.40 | 0.50 | 0.50 |
| Random<br>networks* | Average clustering<br>coefficient±SD | 0.01±0.01 | 0.21±0.03 | 0.20±0.02 | 0.10±0.02 | 0.14±0.02 | 0.08±0.02 | 0.10±0.02 |
|  | Average Harmonic<br>geodesic distance±SD | 4.31±0.25 | 2.40±0.03 | 2.42±0.03 | 2.64±0.03 | 2.51±0.04 | 2.80±0.04 | 2.65±0.04 |

\*Random networks were generated by resetting all of the links of a matching empirical network with the same nodes and links. Data were generated from 100 random runs and SD indicates the standard deviation from the 100 runs.

**Table 2.**
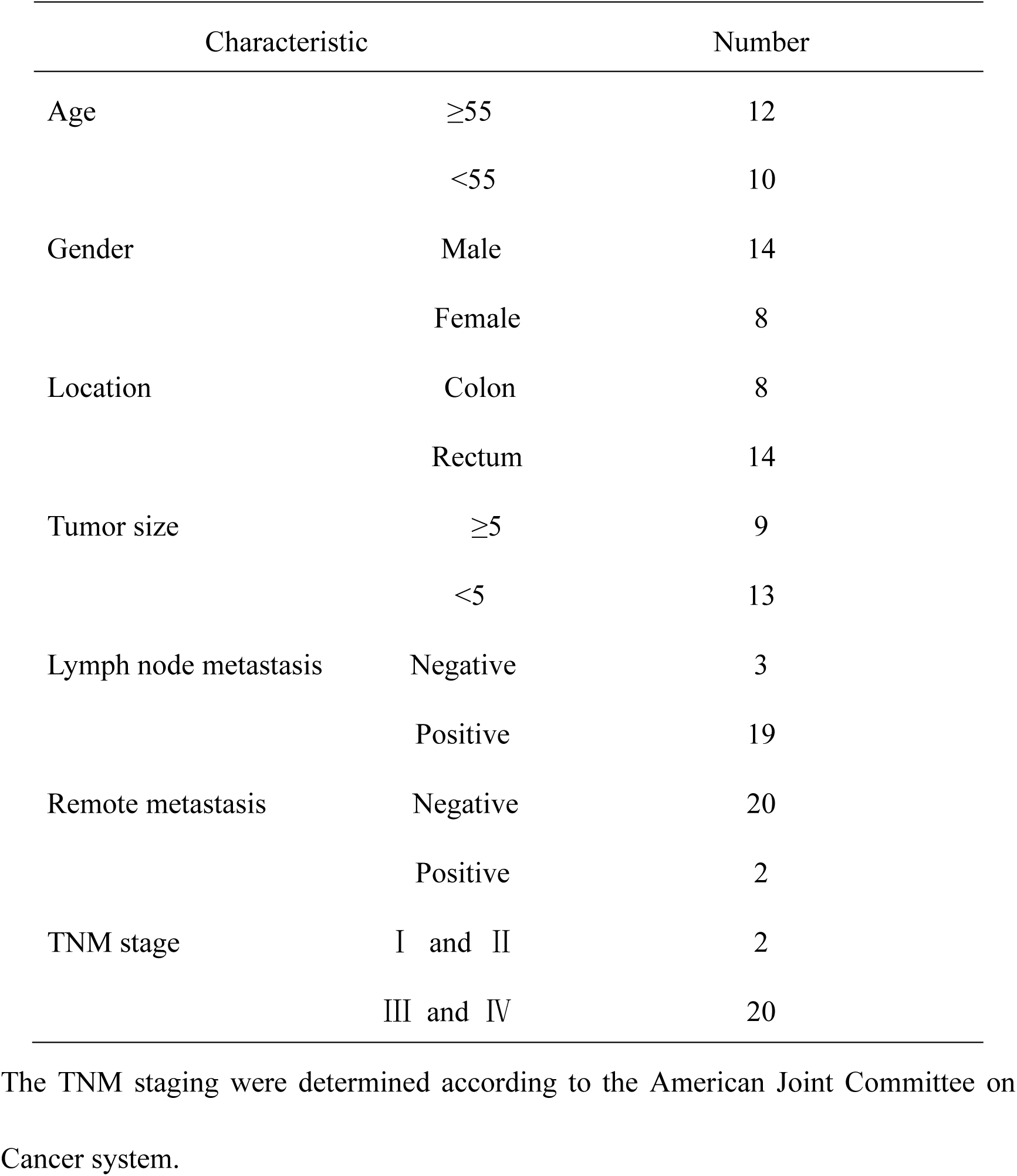
Clinical characteristic of colorectal cancer patients

**Figure 1.**
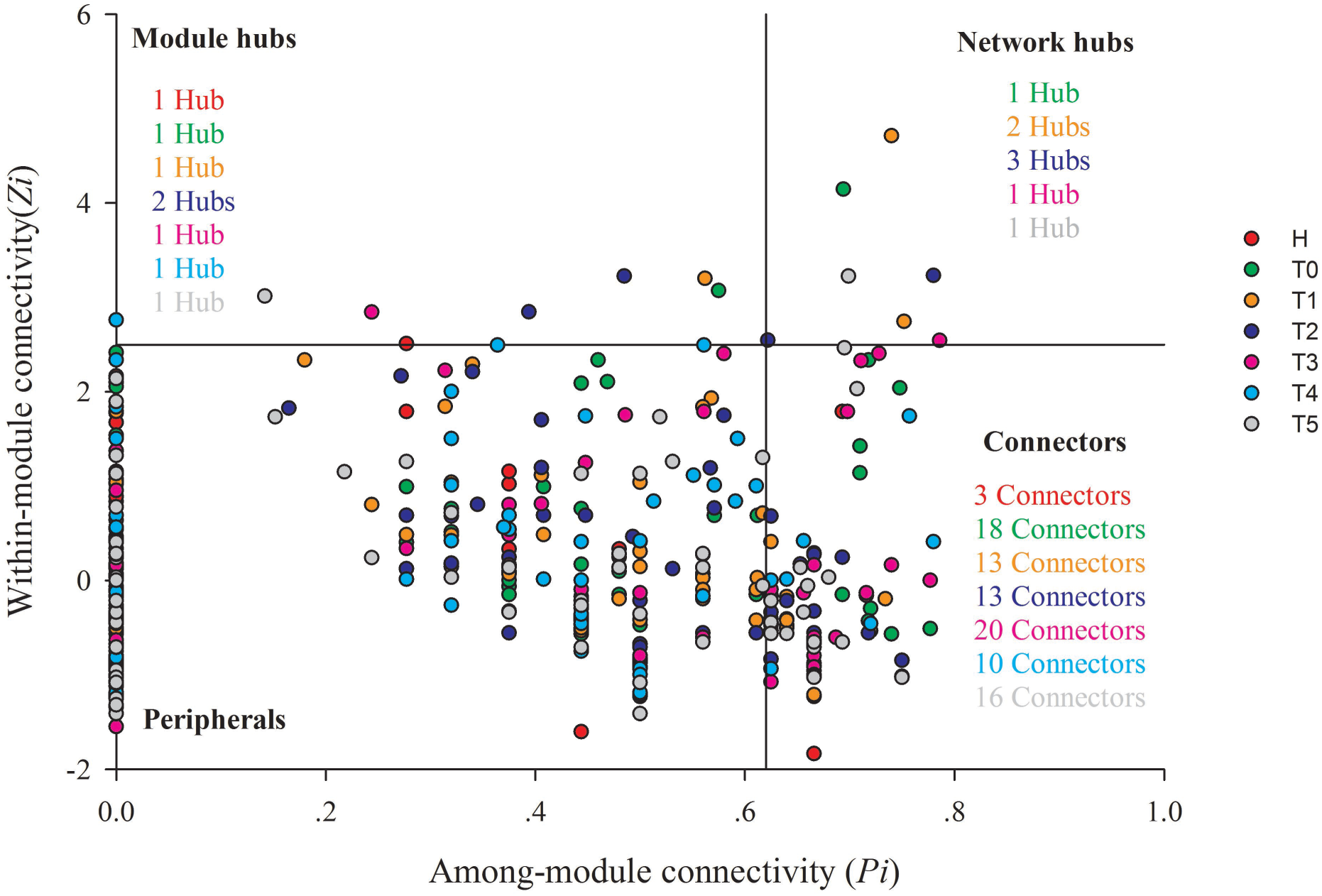
Overview of the networks in healthy individuals and colorectal cancer patients in response to five stages of chemotherapy. Modules are randomly colored at each stage of chemotherapy, and nodes in modules with relative abundance < 1% are in white. Networks represent random matrix theory co-occurrence models derived from more than 19 biological replicates at each stage, where nodes represent OTUs, and links between the nodes indicate significant correlation.

### Distinct networks in colorectal cancer patients and healthy individuals

The intestinal microbial co-occurrence patterns were profoundly different for colorectal cancer patients and healthy individuals networks (Figure 1), which were also proved by multiple network topological properties (Table 1). Healthy assemblages formed larger networks with more nodes than the CRC patients’ networks (Table 1). With the treatment stage going on, the nodes in CRC patients increased ranging from 99 to 103, except for T3 (80). However, the CRC networks contained more links between nodes than healthy networks, which increased the density of connections and created more intricate network patterns (Figure 1 and Table 1). The increased complexity of T networks was also reflected by the shorter harmonic geodesic distances (*HD*) and the increased average degree (*avgK*) (24). However, none of network size and connectivity in CRC patients was close to the healthy individuals. Collectively the above results indicated that the network in CRC patients did not present a regular change with treatment stage increasing, and do not totally close to the healthy network. In addition, we examined the correlation of alpha diversity and network size and connectivity for CRC patients according to multiple univariate diversity metrics. The results showed that there were no significant relationships between them (*P* > 0.05), except that the network size was significantly positively correlated with richness (*P* = 0.025, r = 0.87, Figure S2).

### Modularity in CRC intestinal communities

To identify microbial assemblages that potentially share or interact intestinal niches during the treatment process of CRC patients, we focused on representative networks from CRC patients with five treatment stages. We focused on modules with at least five nodes, and visualized the phylogeny for these modules (Figure 2). Total of 8 modules were detected in the healthy individuals. There were 6, 6, 5, 6, 7, 7 modules, respectively in T0, T1, T2, T3, T4 and T5 network (Table S2). Networks from all CRC patients contained modules with modularity (*M*) values < 0.50 (Table 1). Overall, taxa tended to co-exclude (negative correlations, blue lines) rather than co-occur (positive correlations, gray lines); negative correlations accounted for 47–75% of the potential interactions observed at each treatment stage (Figure 2). The negative correlations in CRC patients decreased from T0 to T5; however, they were still more than that in healthy individuals (45%).

**Figure 2.**
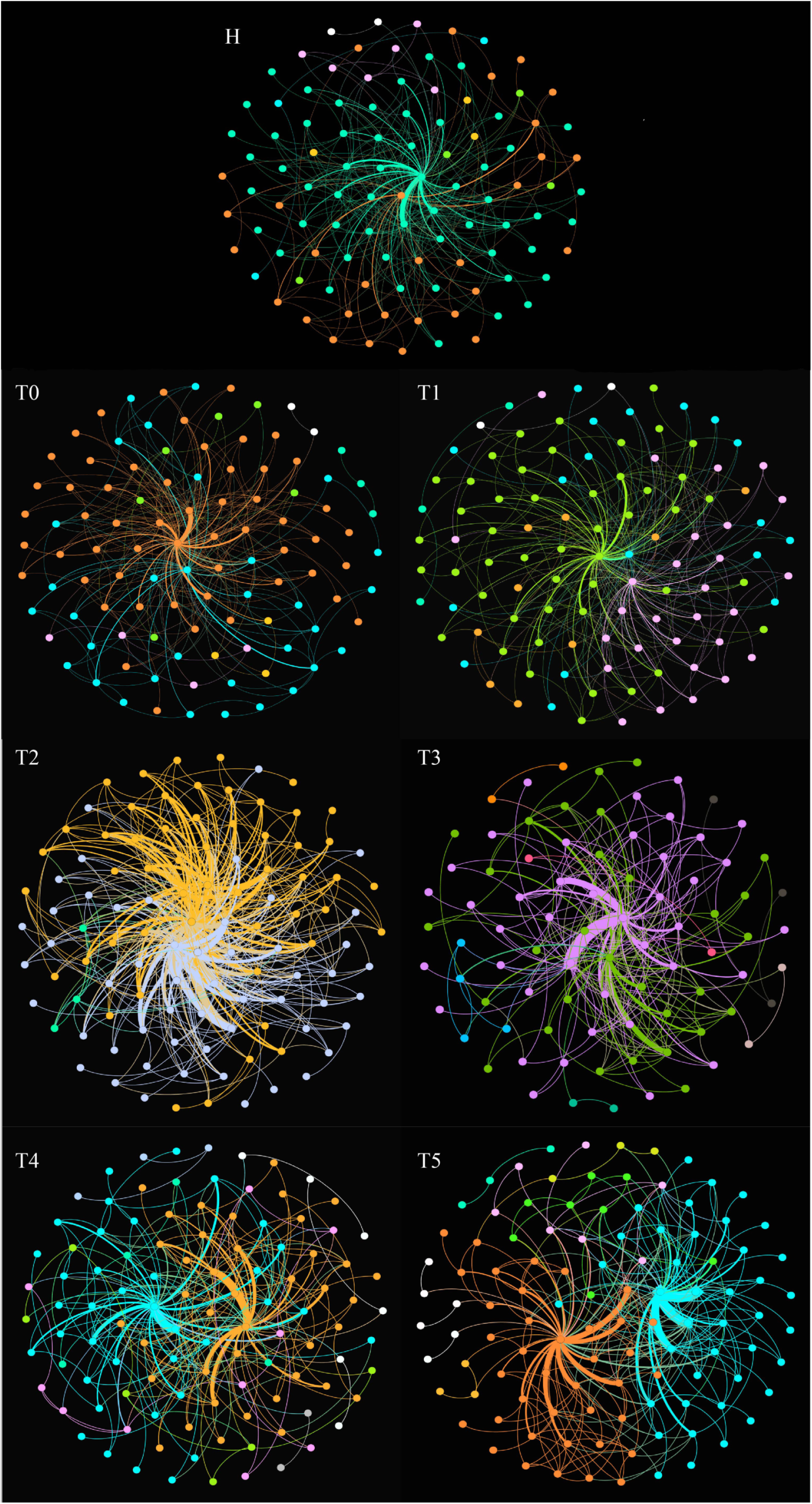
Highly connected modules within intestinal networks of colorectal cancer patients in response to five stages of chemotherapy. Colors of nodes represent different major phyla; pie charts represent the composition of modules with > 1 phyla. A grey link indicates positive relationship between two individual nodes, whereas a blue link indicates negative relationship.

The composition of modules differed within each network and changed over treatment time (Figure 2). *Firmicutes* almost dominated all the modules of each treatment stage of CRC patients. The phylum *Fusobacteria* presented in the module before the treatment (T0), after the first treatment (T1) and the third treatment (T4); and primarily co-exclude with *Firmicutes* or *Bacteroidetes* or *Proteobacteria*, which was supposed to be more relevant to dysbiosis and CRC progression (3, 26).

### Topological roles of taxa in ecological networks of CRC patients

To evaluate possible topological roles of taxa in the networks, we took these nodes into four categories based on the values of among-module connectivity (*Pi*) and within-module connectivity (*Zi*): module hubs, connectors, peripherals and network hubs (24) (Figure 3, Table S3, Table S4 and Table S5). The nodes in each network were mainly peripherals with most of their links inside their modules. Module hubs and connectors have been proposed to be keystone taxa because of their important roles in network topology (24). In this study, the module hubs and connectors were detected in all networks. There were one or two nodes in each treatment stage classified as module hubs in T networks. Total of 7 module hubs identified in T networks originated from a variety of taxonomic groups. The module hubs in T0 and T1 were respectively from the genus of *Klebsiella* and *Fusobacterium* (Table S3), which act pathogen routinely found in human intestine that causes diarrhea and bloodstream infections and markedly increases the rates of treatment failure and death (27). Two taxa from the genus *Bacteroides* and *Atopobium* were classified as module hubs in T2. The T3, T4 and T5 were respectively from the member of *Faecalibacterium*, *Bacteroides* and *Faecalibacterium* (Table S3). Compared with H network, T networks had more connectors during the different treatment stages. The majority of the connectors in each network originated from the *Firmicutes*, and the others were from the *Actinobacteria*, *Bacteroidetes*, *Proteobacteria* and *Candidatus Saccharibacteria* (Table S4).

**Figure 3.**
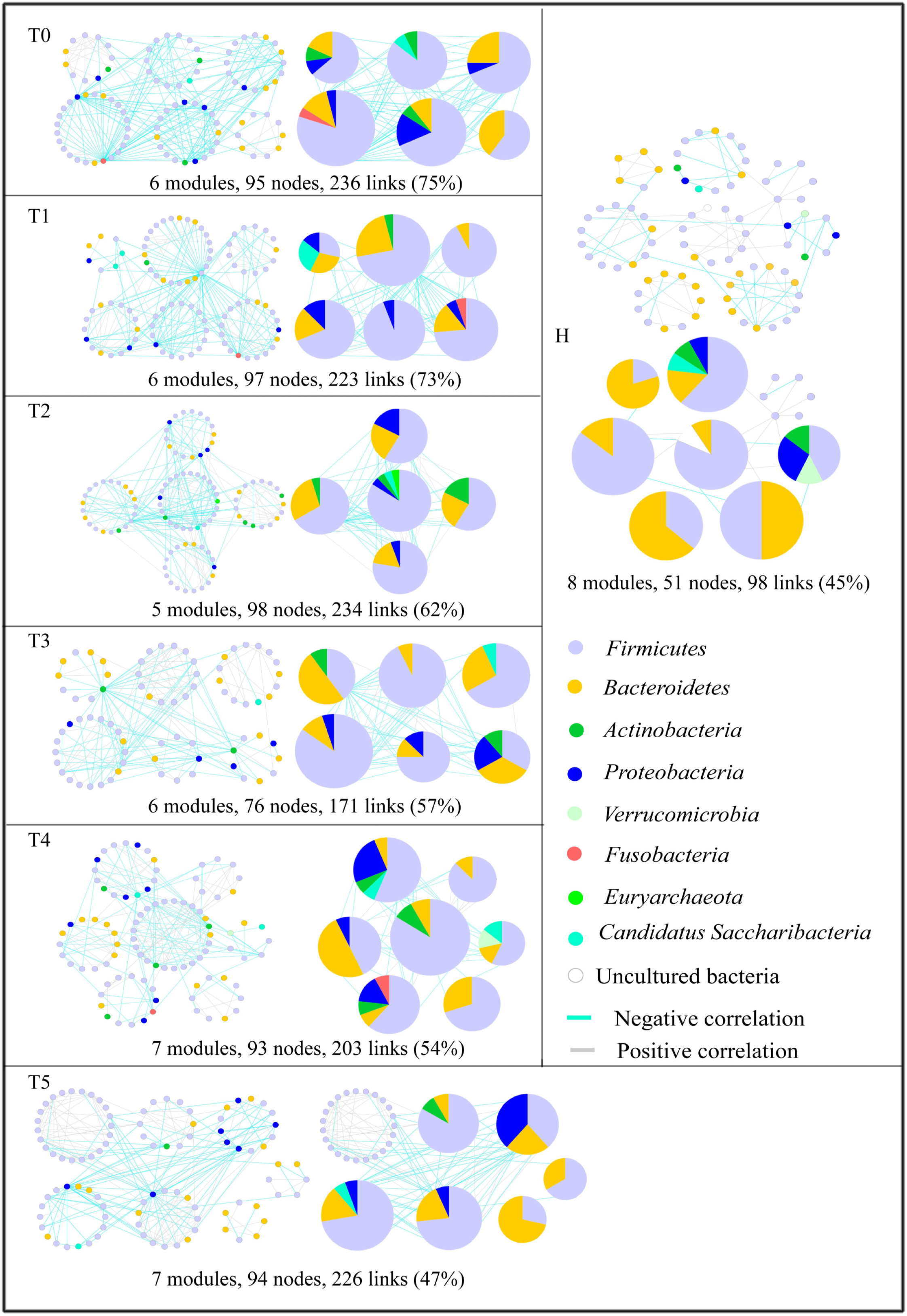
Putative keystone species in the networks of colorectal cancer patients in response to chemotherapy. Different colors represent different stages of chemotherapy, namely red for healthy individuals (H), green for T0, orange T1, blue for T2, pink for T3, water blue for T4, and grey for T5.

In addition, we selected some key OTUs and tumor markers (CA242, CEA, CA199, and CA724) to make Spearman correlation analysis to explore the linkages between microbial correlation networks and clinical chemotherapeutic effects. The results showed that the OTU167, OTU8 and OTU9 were significantly correlated with CEA, CA724 and CA242 (*P* < 0.05, Table S6). The OTU167, OTU8 and OTU9 were from the genus of *Fusobacterium*, *Bacteroides* and *Faecalibacterium*, respectively. These significant OTUs, which were classified into module hubs or network hubs in T0, T1, T2 and T3, were conspicuous in ecological networks, indicated that they were extremely sensitive to chemotherapy.

## Discussions

Chemotherapy remains the mainstay of treatment for patients with CRC, except available surgical debulking (28). In recent years, the changes that occur to the intestinal microbiota are gradually understood in response to cancer chemotherapy. Chemotherapy can damage the mucus layer and disrupt the mucosal barrier in the intestine. Then some intestinal microbiota probably penetrates the lamina propria, leading to life-threatening systemic infections to activate the innate immune system to influence the efficacy of chemotherapy (29, 30). Previous study reported that chemotherapy severely influenced the homeostasis of intestinal mcirobiota (31); in turn, intestinal microbiota modulated the efficacy and toxicity by key mechanisms as translocation, immunomodulation, metabolism, reduced diversity and ecological variation (32). Thus, to optimize current therapeutic strategies, which could help improve the patients’ survival, we compare the intestinal microbiota from the faeces of CRC patients in response to five stages of the chemotherapy based on the molecular ecological network analysis.

Microbes in the intestine are not independent individuals but complex inter-connected ecological communities. Microbes compete for resources or exchange genetic material influencing the microbial composition and host health. Therefore, understanding microbial interactions is important to understand their function. Multiple mechanisms may be responsible for changing network size and complexity in the intestinal microbiota. Chemotherapy distinctly influenced the ecological networks within the intestinal microbiota in CRC patients (T), identified in terms of the average degree (*avgK*), average clustering coeffcient (*avgCC*), harmonic geodesic distance (*HD*) and modularity (*M*). The T networks had their unique ecological network model and varied with different treatment stages (Figure 1). The T networks formed more complex networks than that of healthy individuals, determined by more kinds of phyla in each sub-module (Figure 2). Generally, the OTUs from the same species within a module probably share the same functions (33), and the loss of some OTUs do not disturb the overall function of the module. Accordingly, the less OTU from the same phylum belongs to the same module, the less stable that module would be. Thus, chemotherapy may have key effects on the homeostasis of intestinal microbiota based on the molecular ecological network analysis.

Generally, high alpha diversity is often relevant to health and temporal stability (34). Conversely, low diversity is often observed in the unhealthy intestinal microecosystem contributing to CRC (35). A drastic drop was observed in alpha diversity (Shannon diversity, Simpson diversity, richness and phylogenetic diversity) of intestinal microbiota after the first stage in response to the chemotherapy (Table S1). The similar results were proved by previous study about a steep reduction in alpha diversity of the intestinal microbiota during high-dose chemotherapy for bone marrow transplantation (36). The reduced richness of the intestinal microbiota was also found in obese or IBD patients (37, 38). During the five stage of chemotherapy, T1 had the lowest of Shannon diversity, phylogenetic diversity and richness, except for the Simpson diversity (Table S1). However, the result was inconsistent with the changes in the ecological networks of intestinal microbiota, which was the lowest after the third chemotherapy (T3, Table S2). Pathological disease states might create a state of dysbiosis, and chemotherapy further exaggerated the effect of intestinal microbiota. The intestinal microbiota might adapt to the effect of chemotherapy by increasing the connectivity or the complexity of specie-specie interactions, proved by higher links (Table 1). Therefore, it presented the dissimilarity in the changes of alpha diversity and ecological networks of intestinal microbiota.

The identified modules within the networks probably arise from microbe–microbe interactions in response to shared niches in the intestine. The links in the ecological network could be explained that the co-occur or co-exclude of two nodes, caused by the species performing similar or complementary, and contrary or exclusive functions (39). Study results showed that the links in T modules were distinctly richer than that in H modules (Figure 2), indicated the more complex and compact specie-specie interactions within ecological network. Positive interactions indicate nodes complement or cooperate with each other, while negative interactions signify predation or competition between the taxa. In a relatively healthy intestine, cooperation probably linked with the stable, was found to be the primary interaction in symbiotic intestinal communities (40, 41). Our results showed more negative interaction appeared in T networks than that in H network (Figure 2). However, previous study reported intestinal microbial species were inclined to coexist stably when the system was dominated by competitive interactions (42). Considering the changes of tumor markers before and after the chemotherapy (Supplemental material 1), we supposed that cooperative interactions probably contributed to promote the stability of intestinal microbial community in our study.

Exploring the modular topological roles is a key step in the identification of key OTUs roles in their modules. Topologically, different OTUs play individual roles in the ecological networks (43). Structurally, the loss of peripherals could not influence the functions of ecological networks, while losing the connectors, module hubs or network hubs would result in the deterioration of the entire network (44). In healthy individuals, there were only one module hub, one connector and no network hubs in H network, suggesting the structure of network was relatively stable. There were 18 connectors in T0 network which was distinctly more than that in H network, while the number of connectors did not change much within the ecological networks after five stages of chemotherapy. Therefore, the intestinal microbes were likely to be in a hyperactive state based on the number of network hubs, module hubs and connectors in response to the chemotherapy.

## Conclusions

Ecological molecular network is considered as an effective way to evaluate the intestinal microbiota homeostasis systematically and provides novel insights into the intestinal microbiota in response to the chemotherapy. In this study, we supposed that intestinal microbiota of colorectal cancer patients was still in an unstable state after five stages of chemotherapy based on the ecological molecular network analysis, which might influence the prognosis of colorectal cancer. Therefore, we consider adding probiotics or reducing antibiotic usage to change specie-specie interactions, improve modularity, and enhance the number of generalists to influence intestinal microbial homeostasis. However, further studies need to provide for more evidence to support this idea.

## Materials and methods

### Experiment description

Total of 22 CRC patients from affiliated hospital of Qingdao University (Qingdao, China), aged 34-73 years, were enrolled in our study (Table 1). This study was selected from histopathological diagnosis of primary CRC, newly diagnosed and untreated, and no history of other tumors. All CRC patients were guided by the National Comprehensive Cancer Network Guidelines. The 21 healthy volunteers, aged 27-64 years, were selected as controls. During a routine physical examination, all these participants who used antibiotics within two months, or were regularly using non-steroidal anti-in?ammatory drugs, probiotics or statins before sampling, were excluded from the study. Other exclusions included chronic bowel disease, food allergies, dietary restrictions, and other signs of infections. Based on the national comprehensive cancer network (NCCN) guideline, these CRC patients were performed by standard chemotherapy (45), the following-up samples were obtained from CRC patients before the treatment of every stage, named T0, T1, T2, T3, T4 and T5. Total of 123 fecal samples from the CRC patients were collected for about six times. All these participants were local residents of Qingdao city for more than five years. This study was approved by the Ethics Committee of the Affiliated Hospital of Qingdao University and all study participants signed the informed consent before participation. Fresh fecal samples were collected into 5 ml tubes and immediately stored at −80 °C until the day of analysis.

### DNA extraction, purification, sequencing and data processing

Microbial DNA was extracted from fecal samples by QIAamp Fast DNA Stool Mini Kit as previous reported (46). The freshly extracted DNA was purified by 1% melting point agarose gel followed by phenolchloroform-butanol extraction. The V3-V4 region of the 16S rRNA gene from each sample was amplified by the bacterial universal primers (forward primer: 5’-ACTCCTACGGGRSGCAGCAG-3’ reverse primer: 5’-GGACTACVVGGGTATCTAATC-3’). PCR amplification was performed in a 30μl reaction, containing 15μl 2 × KAPA HiFi Hotstart ReadyMix, 10 ng template DNA, 1μl of each primer (forward and reverse primer), and the rest of the ddH_2_O. The reaction mixtures were operated to a denaturation at 95°C for 1 min, followed by 12 cycles of 98°C for 15 s, 72°C for 10 s, 94°C for 20 s, 65°C for 10 s and 72°C for 10s, followed by 11 cycles of 94°C for 20 s, 58°C for 30 s, 72°C for 30 s, and a final extension at 72°C for 150 s. The PCR amplification products were purified through AxyPrep DNA Gel Extraction Kit (Axygen, USA), eluted in 30μl water, and aliquoted into three PCR tubes. DNA quality and quantity were assessed by the ratios of 260/280 nm and 260/230 nm, and final DNA contents were quantified with a Qubit^®^ dsDNA HS Assay Kit (Invitrogen, USE). Finally, we sequenced bacterial DNA amplicons from each fecal sample based on the Illumina Hiseq 2500.

Raw sequences were separated to samples by barcodes. After quality control of raw data, the clean reads were first sorted by the abundance from big to small, and removed the singletons. Based on the Usearch software, these reads were clustered and removed the chimera based on the standard 97% similarity. The reads of each sample were randomly pumped to extract the corresponding Operational Taxonomic Units (OTU) sequences. Each OTU is considered to represent a species.

### Network construction and analysis

Networks were constructed for intestinal microbiota from CRC patients and healthy individuals based on OTU relative abundances, yielding a total of 7 networks. Covariations were calculated across more than 19 biological replicates to create each network. Only OTUs detected in more than 50% replicate samples were used for network construction. Before the network construction, the random matrix theory (RMT), which is a reliable, sensitive and robust tool for analyzing high-throughput genomics data (47), was used to automatically get the appropriate similarity threshold (St) for identifying modular network and elucidating network interactions. St means the minimal strength of the connections between each pair of nodes (48). Global network properties were characterized according to previous research (24). Modules reflect divergent selection regimes, clusters of phylogenetically and closely related species, habitat heterogeneity and the key unit of species co-evolution (49). Modularity is used to demonstrate a network which is divided into distinct sub-groups naturally (47). Average degree (*avgK*) represents the complexity of the network. Harmonic geodesic distance (*HD*) means the state (close or dispersive) of the nodes in the network. Average clustering coefficient (*avgCC*) is described to that how well a node is linked with its neighbors. The modular topological roles are explained by peripherals, connectors, module hubs and network hubs, which are defined by two parameters, within-module connectivity (*Zi*) and among-module connectivity (*Pi*) (23, 47). In an ecological perspective, peripherals tend to the role of specialists, whereas connectors and module hubs were considered as generalists and network hubs as super-generalists (50).

### Statistical analyses

The R software package (v3.4.1) was used for all statistical analysis. Alpha diversity was calculated using the observed species, phylogenetic diversity, Shannon diversity and Simpson diversity. Linear regression analyses characterized the slopes for alpha diversity and network parameters (nodes or links) in response to the chemotherapy. All analyses were performed based on Molecular Ecological Network Analyses (MENA) Pipeline (http://ieg2.ou.edu/MENA/), and networks were graphed by the software of Cytoscape 2.8.3 (51) and Gephi 0.9.1 (52).

## Declarations

### Ethics approval and consent to participate

The affiliated hospital of Qingdao University Institutional Review Board approved this study, and all study subjects provided informed consent.

### Consent for publication

Not applicable.

### Availability of data and material

Raw sequences have been uploaded into the NCBI Sequence Read Archive data with Accession No.SRP137370 (https://www.ncbi.nlm.nih.gov/sra/SRP137370).

### Authors’ contributions

All authors were involved in the design of the study. Jing Cong analyzed the data. Dong Liu, Na Zhou and Man Jiang collected samples and analyzed the data. All authors interpreted the data. Jing Cong wrote the manuscript. Jingjuan Zhu, Chuantao Zhang, Tianjun Li, Kewei Liu, Helei Hou and Xiaochun Zhang reviewed and revised the manuscript. All authors read and approved the final manuscript.

## Acknowledgements

The authors thank the CRC patients and healthy volunteers for providing the fecal samples that were used in this study. We would also like to thank Amanda Elmore for reviewing and correcting errors and providing feedback on manuscript drafts.

